# Sterols govern membrane susceptibility to saponin-induced lysis

**DOI:** 10.1101/2025.07.17.665205

**Authors:** Malbor Dervishi, Jan Günther, Jinhui Li, Huriye Deniz Uzun, Hans Christian Bruun Hansen, Thomas Gümther-Pomorski, Anja Thoe Fuglsang, Viviana Monje, Søren Bak

## Abstract

Saponins are natural detergents that interact with cellular membranes, causing deterioration leading to membrane disruption. The magnitude of these effects depends on both the saponin structure and target membrane composition, where sterols play a key modulating role in saponin-membrane interaction. We investigated the influence of different sterol classes on saponin-induced membrane lysis. The bioactive, cytotoxic saponin α-hederin induced permeability in membranes containing zoosterol and mycosterol, whereas phytosterol-containing membranes were resistant to lysis *in vitro*. Similarly, in yeast, α-hederin caused significant cell lysis, while in the ergosterol-deficient *erg3Δ* and *pdr18Δ* mutants, cell lysis was minimal. Supplementing phytosterols to yeast provided resistance to α-hederin-induced lysis. Molecular dynamics simulations provide novel mechanistic insights, showing that the efficacy of the activity of α-hederin is proportional to the sterol type in the membrane. Our findings reveal that while zoosterols and mycosterols render membranes vulnerable to bioactive saponins, phytosterols protect membranes from saponin-induced lysis *in vitro*, *in vivo* and *in silico*.

## INTRODUCTION

Sterols are essential biological lipids that play a crucial role in regulating membrane permeability, fluidity, organelle identity, and protein function in eukaryotic organisms^1^. The three major eukaryotic kingdoms; plants, fungi, and animals each evolved distinct sterols derived from the shared intermediate 2,3-oxidosqualene. These sterols include zoosterols such as cholesterol (CHOL) in animals, mycosterols like ergosterol (ERGO) in fungi, and phytosterols, primarily campesterol (CAMP), β-sitosterol (β-SITO), and stigmasterol (STIG) in plants^2^ (Figure 1A).

**Figure 1:**
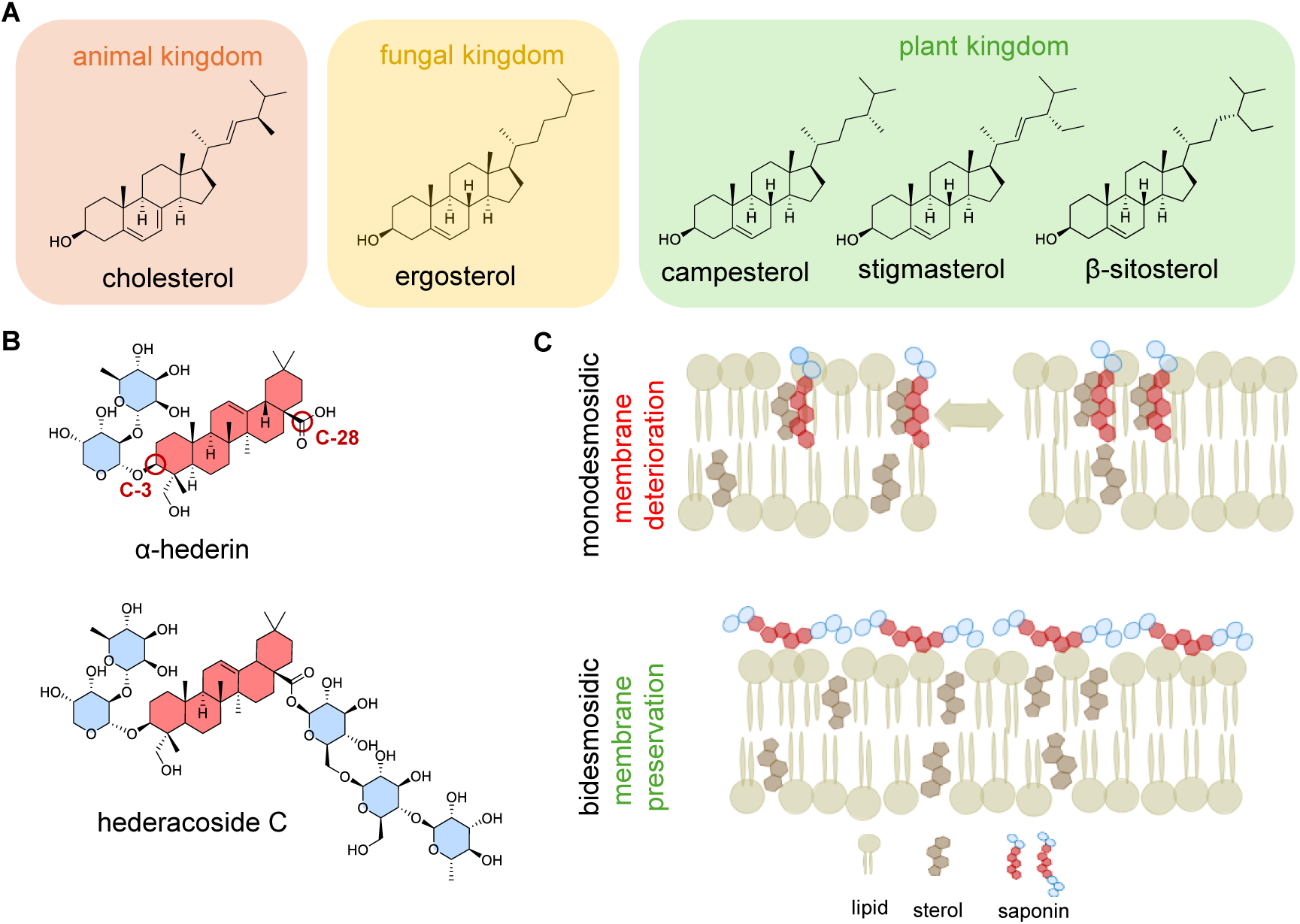
Eukaryotic core sterols determine plant saponin bioactivity. (A) Chemical structures of five different sterols categorized by their primary occurrence in physiology: cholesterol, the predominant sterol in the animal kingdom (orange); ergosterol, characteristic of fungi (yellow); and the phytosterols campesterol, stigmasterol, and β-sitosterol, found in plants (green). These sterols were used in this study. (B) Structures of two hederagenin-derived saponins: monodesmosidic α-hederin (left) and hederacoside C (right), with their core backbone highlighted in red and sugar moieties in blue. Sugar groups are linked at C-3 via an ether bond and/or at C-28 via an ester linkage. (C) Hypothesis illustration of saponin-membrane interactions, comparing monodesmosidic and bidesmosidic saponins. Monodesmosidic saponins readily interact with membrane sterols, leading to membrane disruption, whereas the more hydrophilic bidesmosidic saponins exhibit lower membrane affinity, preserving membrane integrity.

In addition to membrane sterols, plants biosynthesize a vast diversity of specialized triterpenoids originating from the common substrate 2,3-oxidosqualene^3,4^. Among these triterpenoids, saponins constitute a particularly diverse group of naturally occurring amphiphilic, glycosylated defense compounds that typically protect plants against insect pests and pathogens. Structurally, saponins are amphiphilic glycosides composed of a hydrophobic triterpenoid or steroid aglycone (sapogenin) covalently linked to one or more hydrophilic sugar chains. These sugar moieties are typically attached at the C3 and/or C28 positions, giving rise to monodesmosidic (single sugar chain) or bidesmosidic (two sugar chains) forms (Figure 1B). Triterpenoid saponins are widely recognized for their potent antifungal, antibacterial, and antiviral properties^5^ and have attracted growing interest in agriculture and industry as bio-pesticides, pharmaceuticals, vaccine adjuvants, and natural emulsifiers^6–8^. Approximately 100 different triterpene backbones are known, giving rise to over 2000 different saponin structures^9–11^. Found in over 100 plant families^12,13^, most known triterpenoid saponin-producing species belong to the dicotyledons^13–15^.

Despite their bioactive properties, the selective mechanisms by which saponins disintegrate cellular membranes and trigger cell death in non-host organisms remains poorly understood – especially how plants avoid self-damage while biosynthesizing and storing membrane disrupting saponins. It has been hypothesized that saponins insert into membranes via their hydrophobic sapogenin moiety, interact with membrane sterols, and form complexes with sterols and induce membrane curvature, potentially resulting in pore formation, sterol extraction, or disruption of lipid domains^12^ (Figure 1C). It has previously been shown that saponin-induced membrane disruption may depend on interactions with CHOL or ERGO^16^. Yet, the influence of sterol identity on these interactions remains poorly characterized.

In this study, we investigate two structurally related hederagenin-derived saponins: the monodesmosidic saponin α-hederin, that has been shown to be a potent membrane disrupting agent across phylogenetically diverse organisms, and the bidesmosidic derivate of α-hederin, hederacoside C, which is significantly less active ^17^ (Dervishi et al., 2025; under review).

To elucidate the role of membrane sterol composition in modulating saponin susceptibility, we investigate the membrane-disruptive properties of these two selected saponins by examining their interactions with sterols across the three major eukaryotic kingdoms. We use *S. cerevisiae* as an *in vivo* model system, utilizing the wild-type strain (BY4741) and two mutants impaired either in ERGO biosynthesis (*erg3Δ*) or ERGO transport (*pdr18Δ*) to investigate the sterol concentration influence on saponin activity. Under anoxic conditions, yeast cannot synthesize ERGO *de novo* and must incorporate exogenous sterols for membrane integrity and growth^18^. This dependency allows experimental alterations of membrane sterol content. To complement the *in vivo* data, we employed Large Unilamellar Vesicles (LUVs) to quantify saponin-induced membrane permeabilization *in vitro* via calcein release assays^16, 19^. Additionally, *in silico* all-atom molecular dynamics (MD) simulations provide atomic-level insights into saponin-sterol interactions, revealing molecular mechanisms underlying their bioactivity^20–22^.

By integrating experimental and computational approaches, this study advances our understanding of saponin bioactivity, highlighting the role of sterol function in membrane disruption and shedding light on how plants potentially avoid self-toxicity while deploying saponins for defense. These findings provide a mechanistic framework for understanding saponin selectivity and plant self-resistance, offering implications for crop protection and biotechnology.

## METHOD

### Chemicals

The following chemicals were used in the study: α-hederin (Extrasynthesis, France); hederacoside C Analytical standard (Sigma-Aldrich, Germany); propodium iodide (Thermofisher, Austria); 3-(N-morpholino)propanesulfonic acid, MOPS (Sigma-Aldrich, Denmark); polyoxyethylene sorbitan monolaurate, TWEEN^®^ 20 (Roche, Germany); ergosterol (ERGO, >95%), cholesterol (CHOL, >99%), campesterol (CAMP, >65%; ∼35% dihydrobrassicasterol), Stigmasterol STIG >99%, β-sitosterol (β-SITO, >99%), all purchased through Sigma-Aldrich (Germany); 1,2-di-(9*Z*-octadecenoyl)-*sn*-glycero-3-phosphocholine, DOPC (Avanti, USA); Sephadex G-50 (Sigma-Aldrich, USA); bis[N,N−bis(carboxymethyl)aminomethyl]fluorescein, calcein (Sigma-Aldrich, USA); *N*,*O*-bis(trimethylsilyl)trifluoroacetamide, BSTFA (Sigma-Aldrich, Switzerland).

### Yeast Strains and growth

All strains were purchased from Euroscarf (Germany): Wild type BY4741 (*MATa his3Δ1 leu2Δ0 met15Δ0 ura3Δ0*); *erg3Δ* (*BY4741; MATa his3Δ1; leu2Δ0; met15Δ0; ura3Δ0; YLR056w::kanMX4*) and *pdr18Δ* (*BY4741; MATa; his3Δ1; leu2Δ0; met15Δ0; ura3Δ0; YNR070w::kanMX4*).

Yeast strains were cultured overnight at 28 °C and 150 rpm (Innova 44, New Brunswick Scientific, USA) in 20 mL YPD medium supplemented with 20 µg/L carbenicillin, starting at an OD₆₀₀ of 0.2 in 100 mL Erlenmeyer flasks. Cultures typically reached an OD₆₀₀ of ∼10. For each assay, 1mL of culture was transferred to a 2 mL Eppendorf tube and washed three times with buffer A (100 µM MOPS; 1 mM KCl; 2% w/v glucose; pH7, adjusted with Trizma base (Sigma-Aldrich, Denmark). Washing was performed by centrifugation at 14,000 rpm for 2 minutes (Centrifuge 5424, Eppendorf, Germany) to separate the yeast pellet from the supernatant. After the final wash, the pellet was resuspended in Buffer A to an OD₆₀₀ of 0.2^23^. Anaerobic growth was performed under the same conditions using a Microbiology Anaerocult A Mini kit (Millipore, Germany), with 40 μg/L sterol and 0.6% v/v Tween 20 added to the medium^24^. Similar to aerobic growth, each test was carried out by collecting an individual population, and OD₆₀₀ measurements were taken before and after incubation to confirm normal growth.

### Yeast viability assay

Yeast cells were grown overnight in YPD medium, diluted to an OD₆₀₀ of 0.2 in fresh medium, incubated at 28 °C for 3 hours to reach the logarithmic growth phase. Cells were then washed three times with buffer A and resuspended in the same buffer to the original OD₆₀₀. For treatment, 49 μL of the yeast suspension was mixed with 49 μL of propidium iodide (PI) (3 μg/ml dissolved in Buffer A) and 2 μL of the test compound (α-hederin, hederacoside C, or EtOH >96%) in PCR tubes. After 30 minutes of incubation at 28 °C with shaking (150 rpm), 30 μL of each sample was loaded into a chamber of an NC-Slide A2 (100 μm, Chemometec, Denmark). Membrane integrity was assessed using the NucleoCounter NC-3000 (Chemometec, Denmark), which detects PI-bound DNA, indicating membrane permeabilization^25^ (Achilles et al., 2005). Instrument settings were as follows: green channel (excitation 475 nm, emission 560–535 nm), dark field masking; exposure time: 1000; emission filter: Em 675/75; adaptive parameters: maximum number of analyzed cells, and exclusion of aggregates enabled.

### Yeast membrane–Saponin interaction assay

Yeast strains (WT, *erg3Δ*, *pdr18Δ*) were grown aerobically or anaerobically cultured with 40 μg/L sterol and 0.6 % (v/v) Tween 20, as highlighted. After 16 hours, membranes were isolated for LC-MS analysis of saponin interaction. Cultures (1 mL) were washed three times with buffer A, adjusted to OD₆₀₀ = 0.2 in 980 μL buffer A, and exposed (in triplicate) to 20 μL α-hederin or hederacoside C (1 mg/mL) for 15 min. Cells were centrifuged at 10,000 × g (Eppendorf 5424), yielding pellet P1 and supernatant S1. Pellet P1 was resuspended in 5% EtOH, sonicated for 60 min (Branson 5510), and centrifuged again at 10,000 × g to yield pellet P2 and supernatant S2. Supernatant S2 was ultracentrifuged at 100,000 × g (Beckman Optima MAX-XP) to isolate the membrane fraction (Pellet P3) from supernatant S3. Supernatants S1 and S3, and pellets P2 and P3 (resuspended in 85% EtOH), were analyzed by LC-MS^26^.

### Liposome preparation

Large unilamellar vesicles (LUVs) were prepared by extrusion essentially as described^27^. Briefly, DOPC and sterol (9:1 molar ratio) were dissolved in chloroform, mixed, and subjected to three cycles of chloroform evaporation using a rotary evaporator (Rotovap RII, Buchi, Switzerland) to form a uniform dry lipid film. The film was rehydrated with 500 μL loading buffer (62.75 mg calcein in 1 mL buffer A, pH-adjusted with 1N NaOH) and glass beads (212–300 μm), then vortexed for 10 min at maximum intensity (3200 rpm; Vortex-Genie-2, Sigma-Aldrich). Following five freeze–thaw cycles (liquid nitrogen/50 °C water bath), the suspension was extruded 11 times through a 0.2 μm filter using a 1 mL syringe. LUVs were purified by passing the sample twice through a Sephadex G-50 column to remove unencapsulated calcein as described^27^.

### Saponin-Induced Calcein Release Assay

Calcein-loaded LUVs were diluted 1:10 in buffer A, and 180 μL was added per well of a black 96-well plate (Thermo Scientific, Denmark). Saponin (diluted in EtOH) was added (20 μL) to achieve final concentrations of 0–100 μM; EtOH served as a solvent control. After 15 min incubation, 4 μL Triton X-100 was added to lyse vesicles and determine total calcein release. Fluorescence (Ex/Em: 495/515 nm) was recorded in triplicate at three time points: baseline (180 μL), post-saponin (200 μL), and post-lysis (204 μL). Release (%) was calculated using a dilution-corrected formula (1), adapted from Claereboudt et al.^28^:

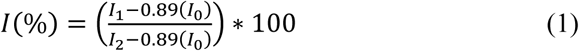

Where I₀, I₁, and I₂ are fluorescence before saponin, after saponin, and after Triton X-100, respectively. A dilution factor of 0.89 was introduced to recalculate the initial intensity before dilution.

### Liposome separation and saponin localization

LUVs (500 μL) were incubated with 20 μM saponin for 15 min at room temperature (∼22 °C). The mixture was then ultracentrifuged at 100,000 × *g* for 60 min (Optima MAX-XP, Beckman Coulter) to separate intact membranes from unbound saponin. Both pellets, membrane fraction (membrane-associated) and supernatant (free saponin) fractions were analyzed by LC-MS.

### Sterol analysis by GC-MS

For sterol analysis, yeast strains (WT, *erg3Δ*, *pdr18Δ*) were grown aerobically; WT was also cultured anaerobically with 40 μg/L sterol and 0.6 % (v/v) Tween 20. Cultures (1ml) were adjusted to OD₆₀₀ = 5.0 before being pelleted and washed three times with buffer A + 2% (v/v) glucose, then lysed in 1 mL 60% KOH containing 10 µM 5α-cholestane (internal standard, >95%, Sigma-Aldrich). Samples were vortexed and incubated at 90°C for 2 hours^29^. Sterols were extracted three times with hexane, pooled, and evaporated at 55°C under nitrogen flow. Samples were subsequently redissolved in 100 μL hexane; 30 μL was mixed with 30 μL BSTFA (Sigma-Aldrich) and derivatized at 60 °C for 1 h.

Derivatized samples were analyzed by GC-MS (Shimadzu GCMS Nexis-2030) using an HP-5MS UI capillary column (30 m × 0.25 mm × 0.25 μm, Agilent). Helium was used as the carrier gas (30 cm/s). A 2 μL sample was injected in splitless mode at 250 °C. The GC temperature program: 60 °C (1 min), ramp to 280 °C over 7 min, then to 310 °C over 36 min, followed by re-equilibration at 60 °C^30^.

### Saponin analysis by LC-MS

Saponins were analyzed on a Dionex UltiMate 3000 UHPLC system (Thermo Fisher Scientific, Germany) coupled to a Bruker Compact qToF-MS with ESI source (Bruker Daltonics, Germany). Separation was achieved on a Kinetex 1.7 μm XB-C18 column using a mobile phase of 0.05% formic acid in water (A) and acetonitrile with 0.05% formic acid (B).The gradient was as follows: 0.0–1.0 min with 5% B; 1.0–2.0 min from 5–30% B; 2.0–14.0 min from 30–70% B; 14.0–15.0 min at 70–100% B; 15.0–16.0 min at 100% B; 16.0–17.0 min back to 5% B; and 17.0–20.0 min at 5% B. Flow rate was 300 μL/min, column temperature was 30°C^31^. MS conditions were as follows: ESI in negative mode, spray voltage −3900 V, dry gas (N₂) flow 8 L/min at 250 °C, nebulizer pressure 2.5 bar, collision energy 10 eV. MS data were collected over *m/z* 50–1400; MS/MS from *m/z* 200–1400 at 3 Hz. Ammonium formate was used for internal calibration via Data Analysis 4.3 (Bruker) as described^31^. Samples were filtered through 0.2 μm Durapore membranes (Millipore) and stored at 4 °C prior to analysis.

### Molecular dynamics simulations: Bilayer setup and Saponin 3D models

Symmetric DOPC:sterol (9:1) bilayers (referred to as membrane-only systems) were built to match the lipid compositions of LUVs using the CHARMM-GUI Membrane Builder^32–34^. Sterols in this study include CHOL, ERGO, CAMP, β-SITO, and STIG. Each bilayer was fully hydrated and neutralized with a 0.15 mol/L KCl ion solution. Membrane-only systems were relaxed using the 6-step CHARMM-GUI protocol for bilayers, followed by a 50 ns production run prior to the insertion of saponin structures^35^. In addition, each membrane-only system was further simulated for an additional 250 ns to characterize the properties of different sterol membranes, which served as controls for the saponin-membrane systems.

Initial 3D coordinates of α-hederin (in protonated or deprotonated states) and hederacoside C were constructed and geometrically optimized using Avogadro^36^ (Hanwell et al., 2012). The structure was parametrized using the CHARMM General Force Field (CGenFF)^37,38^ and solvated using CHARMM-GUI Solution Builder^32,35^. Each saponin system (referred to as saponin-in-water systems) was relaxed using the default minimization and equilibration protocols from CHARMM-GUI Solution Builder. Subsequently, a 150 ns trajectory was run to equilibrate the structure before merging its coordinates in the membrane-only systems. The final 100 ns was used for structural analysis of saponins. All the above membrane-only and saponin-in-water systems were built and run in triplicate to ensure reproducibility and report averages and standard errors accordingly (Table S1).

### Saponin-membrane systems setup

Equilibrated coordinates of the saponins and membranes were merged to examine saponin aggregation, membrane insertion mechanisms, and associated membrane response according to sterols. As before, neutralizing KCl ions were used on each saponin-membrane system to render the simulation box neutral. The simulation of the following starting scenarios: (i) a single saponin molecules positioned 10-15 Å above the membrane; (ii) five saponins of the same type positioned 10-15 Å above the membrane; (iii) five saponin molecules inserted into the membrane. The starting configurations for the third scenario were generated by pulling saponin molecules from the water phase toward membrane tail region using a constant harmonic bias (KAPPA = 10 kJ/mol/nm^2^ for individual saponins) via the MOVINGRESTRAINT function^39,40^ in PLUMED/2.7.1^41–43^. A 10 ns equilibration trajectory was then performed with individual saponins harmonically restrained at the membrane center, defined by the center of mass of the lipid phosphorus atoms in both leaflets. The restraints were finally removed prior to running the unbiased trajectories to examine saponin-membrane interactions (Table S2).

### Simulation settings

All saponin-membrane systems were simulated using the Charmm36m force field^44,45^ and TIP3P water model^46^. As with membrane-only and saponin-in-water systems, each saponin-membrane system simulation was built and run in triplicate to ensure reproducibility and quantify statistical uncertainty. Each replica was simulated for 500 ns, for a total of 42.5 μs of simulation data (Table S3). NPT ensemble at 303.5K and 1 bar using the Nose-Hoover thermostat^47^ and Parinello-Rahman barostat^48^ in GROMACS/2021.5 software package^49^ was utilized. The LINCS algorithm was used to constrain bonds involving hydrogen atoms during the trajectory^50^. Particle Mesh Ewald with a 1.2 nm cutoff was used to evaluate electrostatics^51^, and the van der Waals potential with a force-switching function between 1-1.2nm to account for long-range interactions^52^. Unless otherwise specified, all remaining simulation parameters followed the default settings in GROMACS. Visual Molecular Dynamics (VMD) package^53^ was used to visualize the systems and render snapshots. Trajectory analysis was conducted on VMD, GROMACS, and using the MD Analysis python library^54^.

### Statistical analysis

Group differences were assessed using independent t-tests or one-way ANOVA, as appropriate. Significant ANOVAs were followed by post hoc tests with corrections. Normality and homogeneity of variances were verified prior to analysis. Significance was indicated as p < 0.05 (*), p < 0.01 (**), and p < 0.001 (***).

### EC_50_ Determination

Dose-response data were analyzed using nonlinear regression in R. A four-parameter logistic model (4PL) was fitted using the dose response modeling function from the drc package. The estimated EC₅₀ (half-maximal effective concentration) was fitted using the experimental data using the Levenberg– Marquardt algorithm for parameter estimation. The EC₅₀ value and standard error were extracted directly from the model output using summary() and ED() functions. Confidence intervals were calculated at 95% using profile likelihood methods. Although an initial EC₅₀ value was estimated using concentrations up to 100 µM, the response curve of DOPC:CAMP did not reach a clear plateau (Figure S1; Data S1). This likely resulted in an underestimation of EC₅₀, since the full dose-response relationship was not captured. By extending the concentration range, a more complete sigmoidal curve was obtained, allowing for a more accurate estimation of both the maximum effect (Eₘₐₓ) and the true EC₅₀.

### AI use disclaimer

Artificial intelligence tools, including ChatGPT (GPT-3.5, OpenAI), were used to improve clarity, grammar and structure of the manuscript during initial drafting and revision stages. AI was not used for idea generation, data analysis, scientific interpretation, or reference retrieval. All scientific content, analyses, and conclusions were developed and verified by the authors.

## RESULTS

To assess the role of ergosterol (ERGO) in modulating saponin bioactivity, we used yeast strains differing in ERGO levels (WT, *pdr18Δ*, and *erg3Δ*.). Quantification of ERGO confirmed a significant reduction (40–65%) in both mutant strains compared to WT (Figure 2A, B). Upon treatment with 20 μM of either α-hederin or hederacoside C, cell viability was determined using PI staining. Lysis was significantly higher in WT (75%) than in *pdr18Δ* (43%) and *erg3Δ* (37%) strains, demonstrating an inverse relationship of ERGO abundance and cell viability (Figure 2C; Data S2).

**Figure 2:**
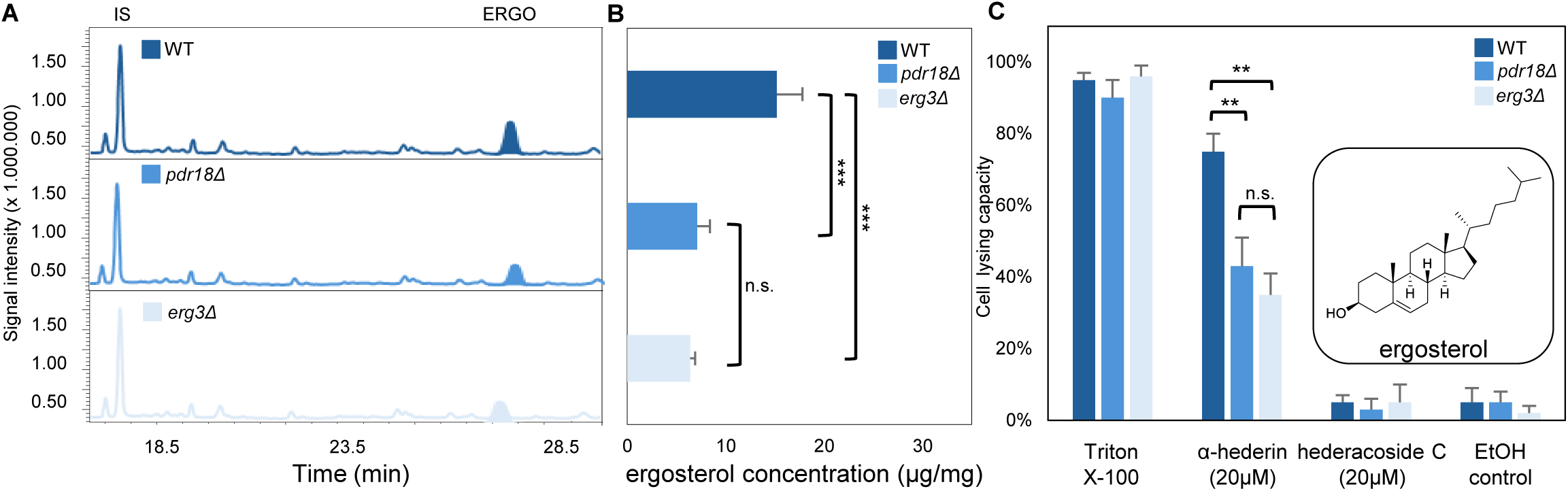
Reduced ergosterol levels confer resistance to α-hederin–induced membrane disruption in yeast. (A) GC-MS analysis of ergosterol content in the wild-type (WT) yeast strain (dark blue) and the mutant strains pdr18Δ (blue) and erg3Δ (light blue). The ergosterol peak at 24.4 min is highlighted for all three strains, highlighting ergosterol reduction of 45–65% in the mutant strains compared to WT, 5-α-cholestane 16.7 min is highlighted as internal standard (IS). (B) Quantification of ergosterol levels in the three yeast strains reveals a significant decrease in membrane-associated ergosterol in *pdr18Δ* and *erg3Δ* mutants relative to WT yeast. (C) Cell lysis capacity of the indicated saponins (20 µM) assessed by PI staining in WT, *pdr18Δ*, and *erg3Δ* strains. Triton X-100 (0.1%) and ethanol (10%) were used as positive and solvent controls, respectively. A reduction in α-hederin-induced cell lysis is observed the ergosterol-deficient mutants. Data represent mean ± SD (n = 5).

These results highlight a direct relationship between ERGO abundance and α-hederin susceptibility, showing the role of ERGO as target for saponin bioactivity in yeast. In contrast, treatment with the structurally distinct bidesmosidic saponin hederacoside C caused no detectable lysis regardless of ERGO abundance, suggesting a significantly different capacity to interact with membrane systems in a lysis-inducing manner.

To investigate sterol-dependent effects on saponin bioactivity, yeast was cultured anaerobically in the presence of individual sterols - cholesterol (CHOL), ergosterol (ERGO), campesterol (CAMP), stigmasterol (STIG), or β-sitosterol (β-SITO) - to enrich the yeast membrane with the respective sterol (Figure 3A). Upon α-hederin exposure, ERGO- and CHOL-supplemented cells showed high levels of lysis (∼75% and 90%, respectively), mimicking the aerobic WT phenotype. Conversely, supplementation with plant-derived sterols conferred resistance to α-hederin: lysis was reduced to 47% (CAMP), 23% (STIG), and 25% (β-SITO) (Figure 3B; Data S1).

**Figure 3:**
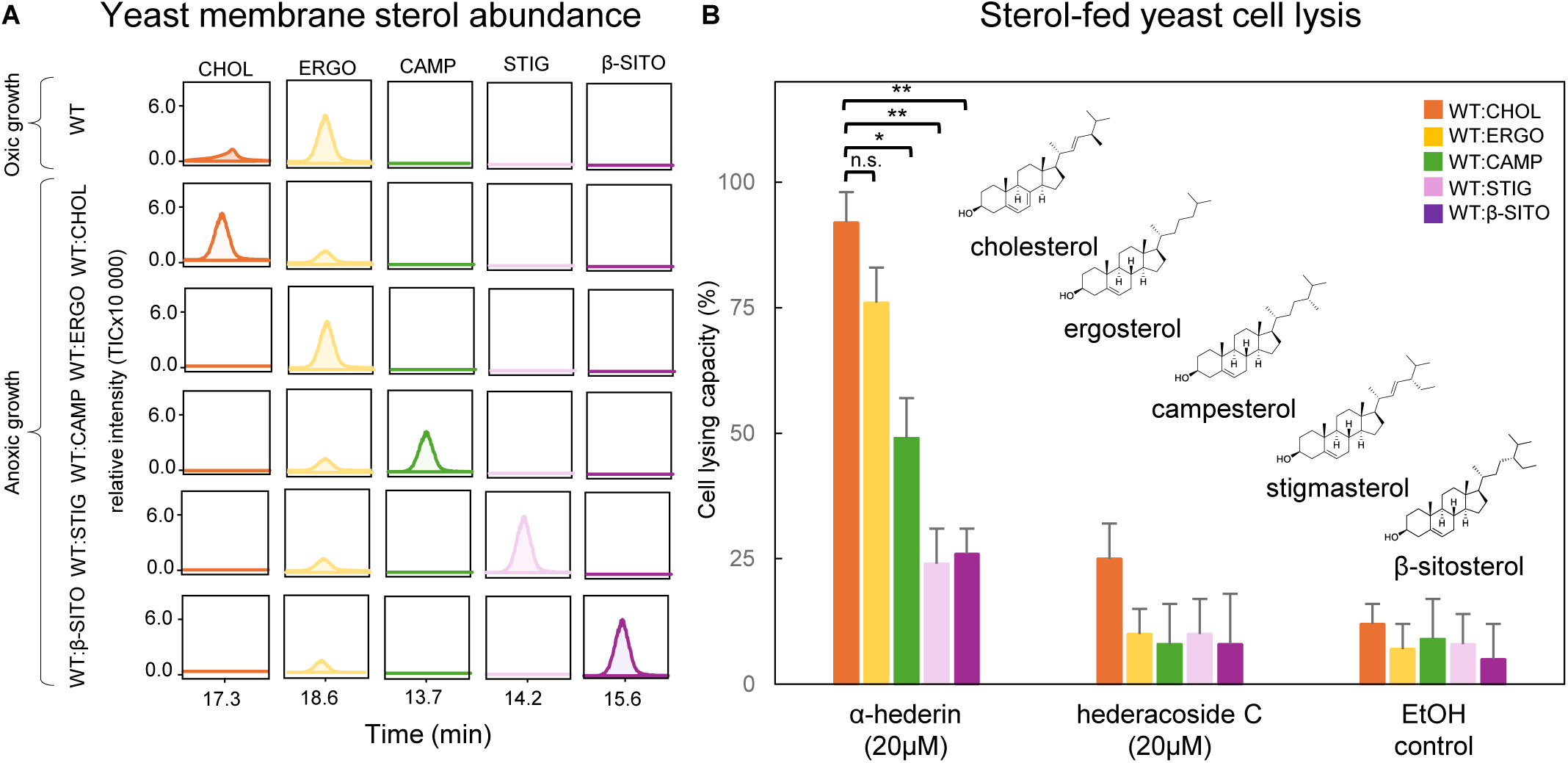
Exogenous sterol supplementation alters ergosterol levels and modulates α-hederin-induced membrane lysis in yeast. (A) GC-MS analysis of sterol composition in wild-type (WT) yeast and anaerobically grown yeast supplemented with exogenous sterols. A reduction in ergosterol production is observed in anaerobically grown strains, while the supplemented sterol is incorporated into the membrane. (B) Cell lysis capacity of saponins and controls in WT strains supplemented with exogenous sterols. A significant decrease in α-hederin activity is observed upon the introduction of phytosterols (n = 5).

These findings demonstrate that membrane sterol composition strongly influences yeast susceptibility to saponin-induced lysis. Membranes enriched with animal or fungal sterols (CHOL or ERGO) are prone to disruption by α-hederin, whereas incorporation of phytosterols significantly enhances resistance. Thus, sterol identity critically modulates both the stabilizing and destabilizing effects of saponins on biological membranes *in vivo*, likely due to differences in sterol packing and membrane organization.

To validate our findings in yeast in a simplified system, we used LUVs composed of DOPC and individual sterols in a 9:1 molar ratio^27^ and treated with increasing concentrations of α-hederin. Calcein-filled LUVs were used to accurately detect membrane lysis. LUV membranes containing either CHOL and ERGO exhibited the highest susceptibility to α-hederin-induced lysis (Figure 4A: Figure S1), with EC₅₀ values of 17.5 ± 2.5 µM and 22.4 ± 3.0 µM for CHOL- and ERGO-containing vesicles, respectively (Figure 4B; Data S2). In contrast, phytosterol-containing LUVs showed increased resistance. CAMP-containing vesicles yielded an EC₅₀ of 54.7 ± 7.3 µM, while STIG and β-SITO did not reach an EC₅₀ within the tested concentration range (0–100 µM). Sterol-free LUVs showed no disruption, confirming that membrane sterols are required for saponin-induced lysis. An upper limit of 100 µM was chosen for α-hederin to remain within the ranges of physiological relevance and to avoid compound precipitation^55^. These findings were supported by MD simulations resulting in higher lysing ratios for membranes containing CHOL or ERGO upon α-hederin exposure (Figure 4C).

**Figure 4:**
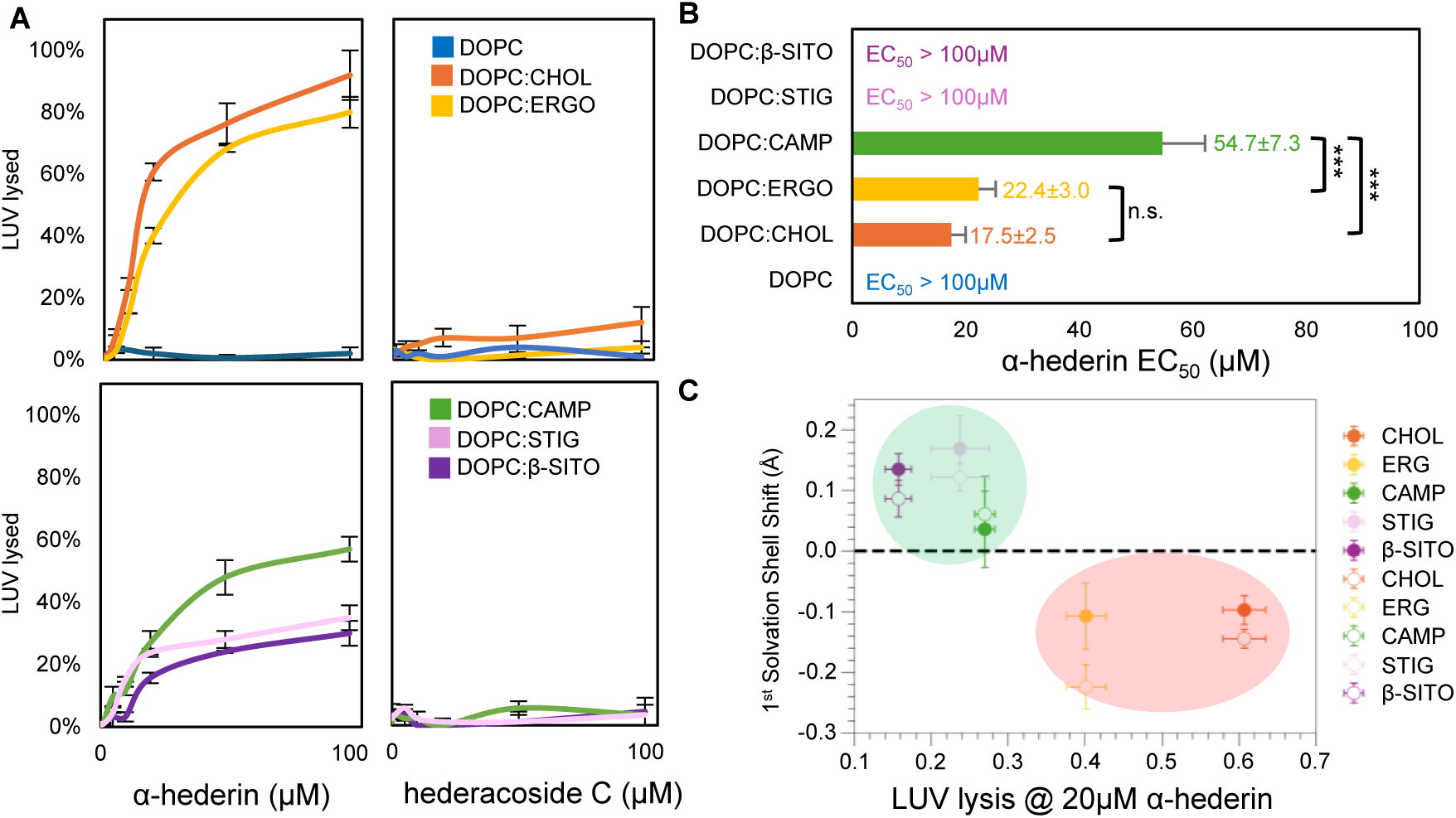
Sterol-dependent membrane disruption by saponins in liposomes correlate with molecular dynamics simulations. (A) Saponin-induced lysis of Large Unilamellar Vesicles (LUVs) by α-hederin (left) and hederacoside C (right) in LUVs containing different sterols (n = 3, mean ± standard deviation) divided in two groups, lipids with no sterol (blue), CHOL (orange), or ERGO (yellow) and group with phytosterols CAMP (green), STIG (pink) and β-SITO (purple). (B) EC_50_ values of LUVs with different sterols, with standard deviation. No EC_50_ values were obtained for DOPC, DOPC:stigmasterol (STIG), and DOPC:β-sitosterol (β-SITO) conditions. (C) Correlation between the shift in the 1st RDF solvation shell from simulation and the lysing ratio of α-hederin (20μM) on DOPC/sterol LUVs *in vitro*. Shift values are calculated as the difference in 1st RDF solvation shell positions between α-hederin-containing and membrane-only systems. Solid markers represent neutral α-hederin-bilayer systems; hollow markers represent deprotonated α-hederin-bilayer systems. The green-shaded region corresponds to a positive shift and lower lysing ratio, and the red-shaded region (including cholesterol and ergosterol) corresponds to a negative shift and higher lysing ratio.

These results demonstrate that the membrane-lytic activity of saponins *in vivo* is extendable to extremely simplified membrane systems *in vitro* and *in silico*, supporting the observation that sterol identity influences saponin-induced lysis.

The two saponins studied - α-hederin and hederacoside C - are representative examples of mono- and bi-desmosidic saponins, where monodesmosidic saponins are generally more bioactive^8^. To determine whether this difference in bioactivity arises from stronger membrane association or higher sterol affinity, we analyzed the distribution of each saponin in membrane and supernatant fractions. In yeast, α-hederin predominantly accumulated in the membrane fraction, especially in ERGO-rich cells. In contrast, when yeast was grown anaerobically with phytosterol supplementation membrane association of α-hederin decreased in ERGO-deficient mutants (Figure 5A). While the inactive hederacoside C was consistently dissociated from membranes, aligning with its minimal lytic potential.

**Figure 5:**
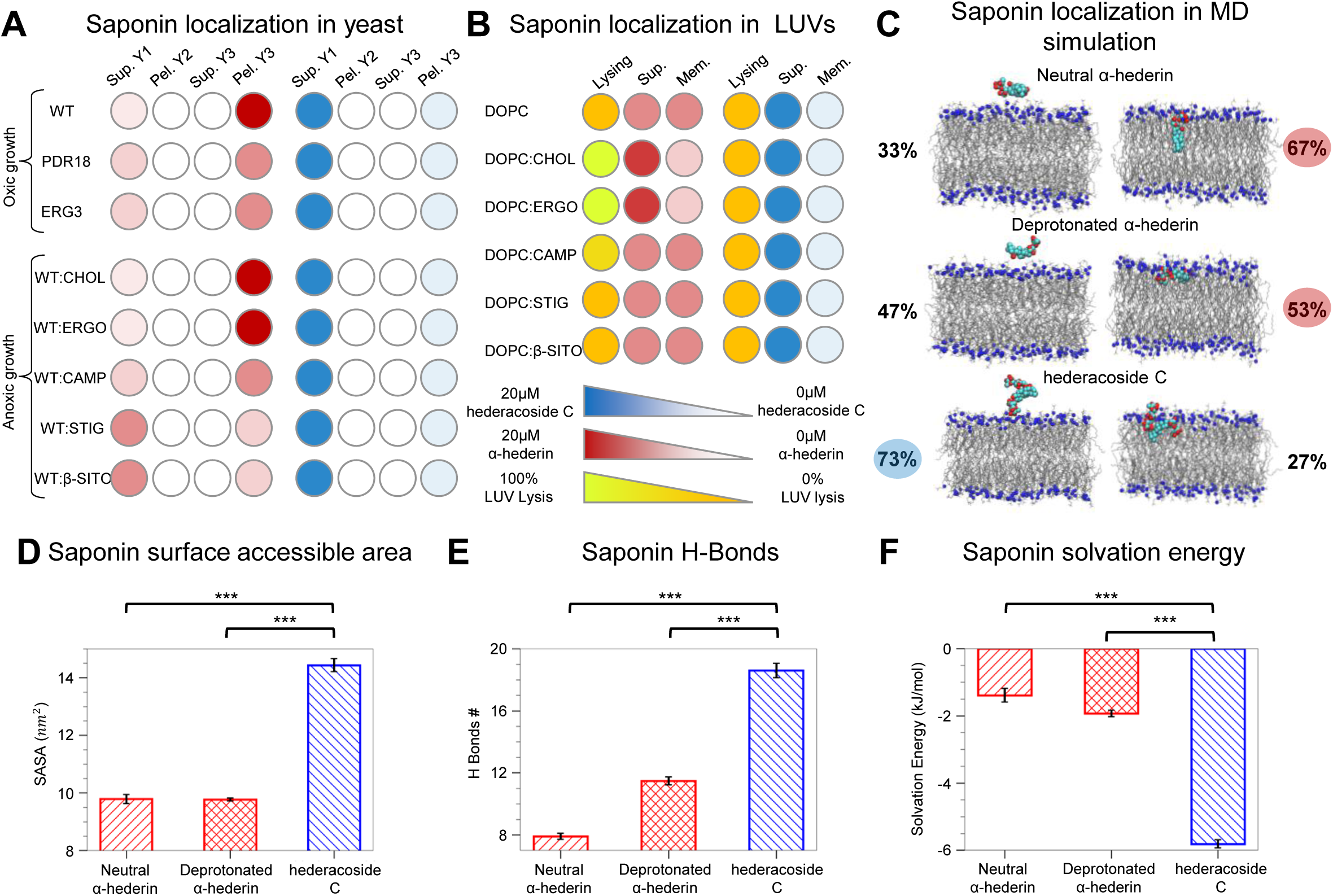
Saponin localization resembles bioactivity upon membrane exposure *in vivo*, *in vitro* and *in silico*. (A) Relative localization of saponins α-hederin (red, left) and hederacoside C (blue, right) in different collected fractions: supernatant 1, pellet 2, supernatant 3, and pellet 3 (membrane fraction) for yeast assay. (B) Lysing levels after saponin exposure together with supernatant and pellet fraction for LUV membranes. Hederacoside C is predominantly found in supernatant for both yeast and LUV assays, suggesting minimal membrane interaction, while α-hederin is enriched in the membrane fraction, indicating potential sterol association. (C) Representative bound and unbound configurations for neutral α-hederin (top), deprotonated α-hederin (middle), and hederacoside C (bottom). Percentages indicate the proportion of systems in which the corresponding event was observed relative to the total number of simulated systems (also summarized in Table S3). (D) Solvent-accessible surface area (SASA), (E) number of hydrogen bonds with water, and (F) solvation energy for neutral α-hederin, deprotonated α-hederin, and hederacoside C. All measurements were obtained from saponin-in-water simulation systems run in triplicate; the error bars represent the standard error across the replicas.

In the LUV experiments, both saponins were detected in the membrane and supernatant fractions. However, CHOL- and ERGO-containing LUVs showed detectable membrane-bound α-hederin, although a significant fraction remained in the supernatant – likely reflecting LUV lysis and soluble membrane fragments. Hederacoside C consistently remained in the supernatant across all LUV compositions, consistent with its minimal membrane-disruptive activity (Figure 5B).

Molecular dynamics (MD) simulations, which showed that approximately 60% of both protonated and deprotonated α-hederin molecules interact with DOPC:CHOL membranes, whereas only 27% of hederacoside C interacts with the same membrane system (Figure 5C). Further analysis of saponin surface accessible area, number of H-bonds and saponin solvation energy of the tested saponins provide further insights into the phase preference of the saponins due to their physicochemical properties highlighting a stronger preference of α-hederin for the membrane bilayer than hederacoside C (Figure 5D-F).

Taken together, these findings suggest that stronger saponin–membrane interactions are critical for the higher bioactivity of α-hederin in comparison to the less active hederacoside C.

To gain atomic-level mechanistic insights of saponin chemical characteristics and interactions with membrane sterols, MD simulations of saponins in water were first used to determine the structural properties of α-hederin and hederacoside C in aqueous solution. Considering the possible ionization of α-hederin and the fixed-charge additive force field used in the simulation, separate simulations were conducted for both the neutral (protonated) and deprotonated α-hederin^37^. A higher solvent accessible surface area (SASA) for hederacoside C compared to either form of α-hederin due to the hydrophilic nature of the sugar moieties in hederacoside C (Figure 5C-E). These features reflect the higher hydrophilicity and provide insight into the stronger preference for the aqueous phase of hederacoside C over α-hederin.

Indeed, membrane simulations revealed fast interactions of α-hederin with the membrane surface, within the first few nanoseconds of simulation. As the simulations progressed, binding and insertion modes differed among different saponin molecules as shown by the z-component distance between the center-of-mass (COM) of the saponins and the membrane (Movie S1). Protonated and deprotonated α-hederin bind to the bilayer in different orientations and insertion depths, while hederacoside C remains in the water phase interacting with the membrane surface intermittently in most cases (Figure 5B). Saponin-membrane interaction events across all MD simulations reveal a lower membrane association of hederacoside C compared to α-hederin (Figure 5B; Table S3). In addition, five hederacoside C molecules were pulled into the tail region of a DOPC:CHOL bilayer using external harmonic potentials. After equilibration, the restraints were removed, and an unbiased simulation was performed. When initially forced into the membrane, one hederacoside C molecule gradually dissociated and returned to the water phase due to its strong hydrophilicity, which is opposed to the higher hydrophobicity of α-hederin (Movie S2). These findings provide detailed mechanistic insight into the distinct chemical properties of saponins and their interaction with membrane systems, further validating the experimental observations of differential membrane affinity. Therefore, the more hydrophobic character of α-hederin favors membrane deterioration and disruption, while the hydrophilic extended sugar chains of hederacoside C precludes stable bilayer interaction.

To dissect how different sterols modulate membrane susceptibility to α-hederin, we performed MD simulations with multiple α-hederin molecules interacting with DOPC:sterol bilayers. In systems with a 200:5 lipid/saponin ratio, both neutral and deprotonated α-hederin form stable aggregates that largely remain in the water phase with limited bilayer interaction, as evidenced by early clustering events observed in the trajectories (Figure S2A). However once inserted into the membrane layer, molecules distribute and rarely leave the lipid bilayer (Figure S2B). To obtain molecular insights into how sterol structures affect the α-hederin lytic activity, five saponin molecules were placed directly into the bilayer center at initiation. After removing the restraints to position the saponins at the bilayer center, protonated molecules remained closer to this location, while deprotonated α-hederin drifted to the membrane surface, remaining below the P-atom layer (Figure S2C and D). Top-view snapshots and cumulative distribution maps show sterol distributions in membrane-only systems and those containing five bound neutral or deprotonated α-hederin molecules over the final 100 ns of simulation (Figure 6A). This pattern was quantified using radial distribution function (RDF) of the relative distribution of sterol hydroxyl oxygen atoms (Figure S2) to quantify the membrane response to the insertion of five α-hederin molecules. The first solvation shell of CHOL and ERGO exhibited a leftward shift, whereas phytosterols exhibit either no shift or a rightward shift upon saponin insertion (Figure S3; Figure 6B).

**Figure 6:**
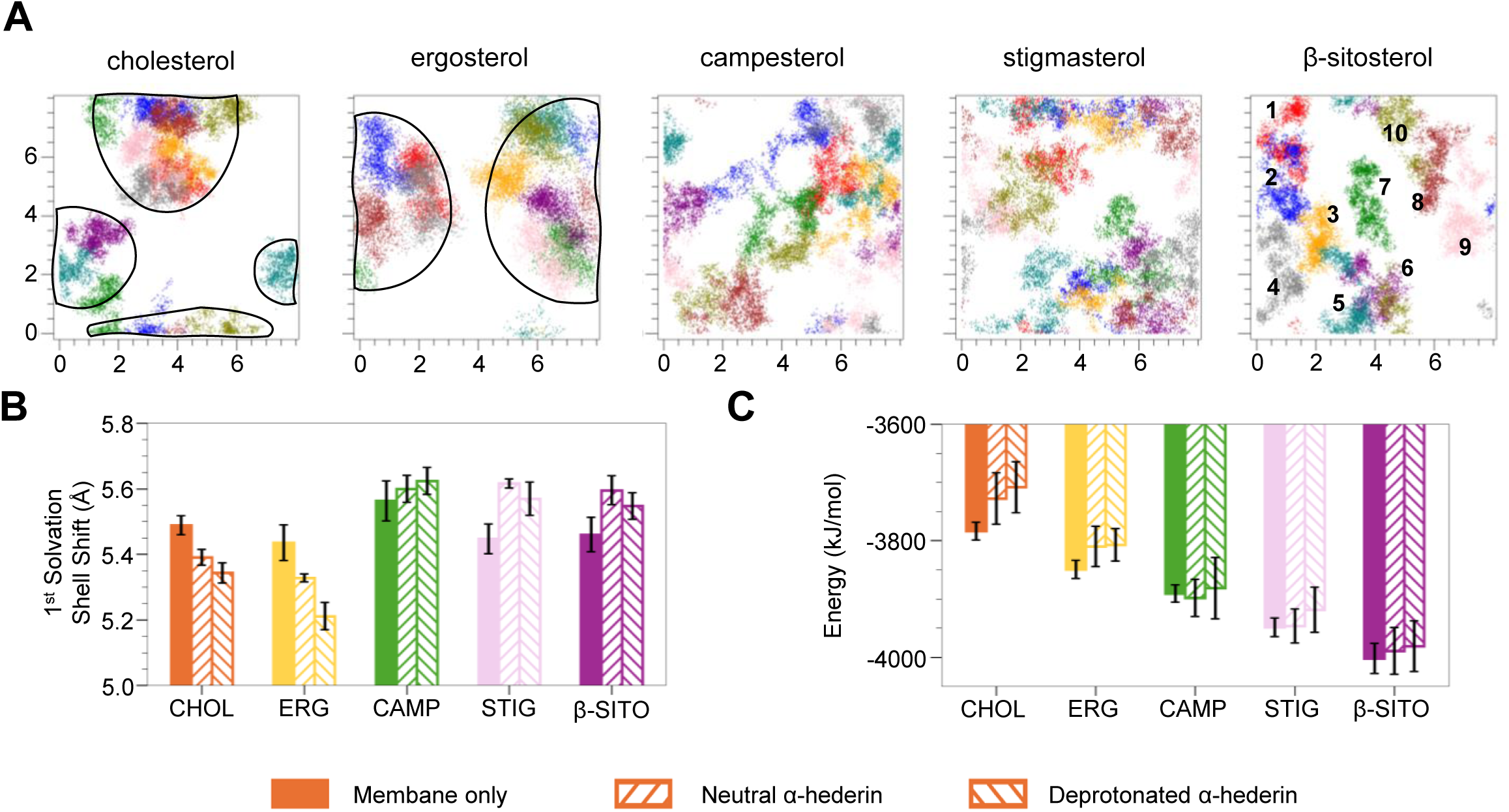
Molecular modeling of membrane-saponin interaction provides mechanistic insights into saponin bioactivity. Representative top-view snapshots and corresponding cumulative sterol distribution maps from the last 100 ns of simulations for membranes with five bound deprotonated α-hederin molecules. Colorful scatter points in the distribution map represent the positions of the individual sterol hydroxyl oxygen atoms. Individual cholesterol molecules are labeled in the last column as reference. We observe a higher aggregation of the animal and fungi sterols (black line) compared to the phytosterols. (B) Position of the 1st solvation shell in 2D radial distribution function (RDFs) for sterol hydroxyl oxygen atoms in membrane-only simulation systems (solid bars) and membranes containing five neutral (left-diagonal filled bars) or deprotonated α-hederin molecules (right-diagonal filled bars). (C) Corresponding non-bonded interaction potential energies between DOPC and sterols.

Importantly, these structural changes correlated with experimental observations that membranes with greater sterol clustering (CHOL and ERGO) were more prone to lysis by α-hederin, while those with dispersed sterols (CAMP, STIG and β-SITO) were more resistant (Figure 6). Thus, sterol clustering appears to increase local instability and thereby significantly destabilizes the membrane system in the presence of α-hederin, potentially promoting pore formation and membrane lysis.

To elucidate the mechanism behind different sterol distribution preferences, the DOPC-sterol nonbonded interaction energy was computed for each sterol species to examine their interaction strengths with DOPC (Figure 6C; Figure S4). A decreasing trend in interaction strength is observed from β-SITO to CHOL, which contrasts with the increasing lysis (Figure 4A). This suggests that sterols with weaker lipid interactions (like CHOL and ERGO) are more susceptible to saponin-induced disruption.

These results provide valuable molecular insights into the sterol-dependent membrane interactions of saponins, a finding that is consistent with experimental observations and highlights the importance of sterol composition in modulating membrane disruption – validated by *in vivo*, *in vitro* and *in silico* approaches.

## DISCUSSION

Saponin bioactivity involves membrane deterioration and perforation, disrupting the chemiosmotic barrier, causing leakage of cellular components from the cell, causing metabolic breakdown and ultimately death. However, the precise molecular mechanisms of saponin-induced membrane lysis remain unclear. We selected two saponins from *Hedera helix,* the monodesmosidic α-hederin and the bidesmosidic hederacoside C, to investigate membrane lysis mechanisms using a combination of *in vivo*, *in vitro*, and *in silico* analyses. Our data demonstrates that variations in sterol membrane composition significantly influence the membrane-lytic activity of plant triterpenoid saponins, with fungal and animal sterols enhancing susceptibility while phytosterols provide resistance.

To test whether α-hederin specifically interacts with sterols in yeast membranes, two yeast mutants (*erg3Δ* and *pdr18Δ*) impaired in ERGO homeostasis were studied. Although these mutants have been shown to be more resistant to various bioactive compounds^56^, our results demonstrate that both *erg3Δ* and *pdr18Δ* mutants—characterized by impaired ERGO accumulation in their membranes—are more susceptible to saponins previously considered highly bioactive^57^. This suggests that ERGO content may correlate with saponin sensitivity in wild-type yeast. Additionally, under anaerobic conditions, wild-type yeast exhibited sterol auxotrophy, and growth was restored by supplementation with various exogenous sterols. Specifically, supplementation with the eukaryotic crown sterols—mycosterol (ERGO) or zoosterol (CHOL)—rendered yeast susceptible to α-hederin, whereas phytosterols reduced the lytic activity of α-hederin. These results indicate that sterols mediate saponin interactions and membrane disruption in a sterol-specific manner. However, the complex biomolecular composition of yeast may obscure saponin-sterol interactions, masking the observed cell lysis. Johnston et al. found that the *erg3Δ* strain is more tolerant to the anti-inflammatory saponin escin^57^. Additionally, simultaneous addition of ERGO and escin to *erg3Δ* strain prevents escin-mediated growth inhibition, while addition of phytosterols did not show the same protective effects^57,58^. The protective role of phytosterols against α-hederin supports the hypothesis that saponin-induced lysis requires compatible sterols, while incompatible sterols like phytosterols reduce susceptibility. Our findings imply that the co-evolution of saponin biosynthesis and membrane sterol composition in plants has resulted in a refined chemical self-defense system, in which phytosterols function not only as structural components of plant membranes but also as protective agents that mitigate the toxicity of the plant’s own saponins.

The lytic activity of α-hederin varies with sterol type, where plant-derived sterols reduce membrane disruption more effectively than CHOL or ERGO, as demonstrated by experiments using LUVs with varied sterol compositions. This finding agrees with previous studies showing that α-hederin is most effective at lysing LUVs containing CHOL, while those lacking sterols remain intact^16^. Lysis was not observed in LUVs composed of DOPC alone, whereas LUVs containing CHOL or ERGO exhibited EC_50_ values of 17.5 µM and 22.4 µM, respectively, confirming the essential role of sterols in saponin-mediated membrane disruption^59^. LUVs incorporating CAMP displayed higher EC_50_ values in response to α-hederin, compared to those containing CHOL or ERGO. For LUVs with β-SITO or STIG, EC_50_ values could not be determined within the tested concentration range, suggesting that sterol identity significantly influences both membrane stability and resistance to saponin interaction. CHOL appears to enhance saponin activity by promoting sterol-packing defects and facilitating pore formation, whereas phytosterols such as β-SITO, STIG, and CAMP likely stabilize membrane packing and mitigate permeability caused by saponin-induced fluctuation (Figure 6A).

The molecular dynamics simulations revealed that α-hederin causes aggregation of CHOL and ERGO in DOPC membranes. This redistribution potentially promotes the formation of lipid microdomains that favor pore formation^60^. Such effects were not observed with other sterols, including Δ7-sterol, highlighting the sterol-specific nature of membrane susceptibility^28^. These findings support a model where sterol composition shapes the membrane response to saponins. Similar mechanisms in organisms like sea cucumbers (*Cucumaria frondosa*), which modify sterols to protect against their own bioactive saponins, suggest broader biological relevance^61^. The species-specific role of sterols in modulating saponin toxicity is emphasized by these findings, providing valuable insights into the sterol-specific mechanisms underlying saponin-induced membrane disruption.

The compatibility between saponins and sterols plays a critical role in determining saponin toxicity that implies that the evolution of alternative sterols could be a key mechanism for saponin resistance in higher eukaryotes. We substantiated our investigations of the interactions between saponins and membranes by *in silico* modeling. Monodesmosidic saponins, such as α-hederin, which contain a single glycosidic chain, can integrate into lipid bilayers in both neutral and deprotonated states due to the more accessible hydrophobic aglycone. During membrane insertion, the deprotonated acid moiety of saponins remains mobile in the aqueous phase and can associate with the membrane interface without requiring prior protonation, as supported by recent findings^62^. While protonation of the carboxylic acid at C28 of the hederagenin backbone may occur upon deeper insertion into the membrane, potentially enhancing hydrophobicity and membrane penetration. In contrast, bidesmosidic saponins, which possess two glycosidic moieties, are sterically hindered from entering the cell^63,64^. Computational analysis of the structural characteristics of both saponin types, including solvent-accessible surface area (SASA), hydrogen bond formation, and solvation energy, revealed that the additional sugar moieties of bidesmosidic saponins (e.g., hederacoside C) lead to a stronger hydrophobicity. As a result, bidesmosidic saponins exhibit minimal stable binding to membranes compared to monodesmosidic saponins, which leads to reduced membrane destabilization and lytic activity. However, under natural conditions, these bidesmosidic saponins may serve as storage forms of the more active monodesmosidic saponins, that can be rapidly released by plant β-glucosidase upon plant tissue rupture, as shown in barrel clover (*Medicago truncatula*), or by glucosidases of target organism acting as a ‘saponin bomb’^65^. The majority of hederacoside C exposed to the yeast and LUV populations remained in the aqueous phase, indicating minimal interaction with the membrane. Conversely, α-hederin exhibited strain-dependent distribution, localizing predominantly within the purified membrane fraction—an indication of its membrane association. Taken together, our *in vivo, in vitro,* and *in silico* results show that α-hederin disrupts membranes via a synergistic action of its hydrophobic aglycone and glycosidic chain, which together destabilize lipid bilayers by perturbating lipid-lipid interactions, inducing membrane lateral packing changes, and ultimately promoting pore formation and lysis.

This study demonstrates that saponin-induced membrane lysis is governed by both saponin structure and membrane sterol composition. We propose a two-component model of saponin bioactivity, in which the molecular structure of the saponin and the physicochemical properties of the sterols together determine lytic potential (Figure 7). While saponin insertion into membranes appears consistently across different sterol types, the extent of membrane disruption varies, with higher lysis observed in CHOL- and ERGO-containing membranes. This suggests that sterol identity influences the degree of membrane destabilization rather than saponin incorporation itself. Such a mechanism offers a compelling explanation for how saponin-producing plants avoid self-toxicity by incorporating non-compatible sterols like phytosterols into their own membranes. Overall, this model helps explain saponin selectivity and supports the idea that the evolutionary diversification of saponins is closely linked to the diversity of membrane sterols across species^28,61^.

**Figure 7:**
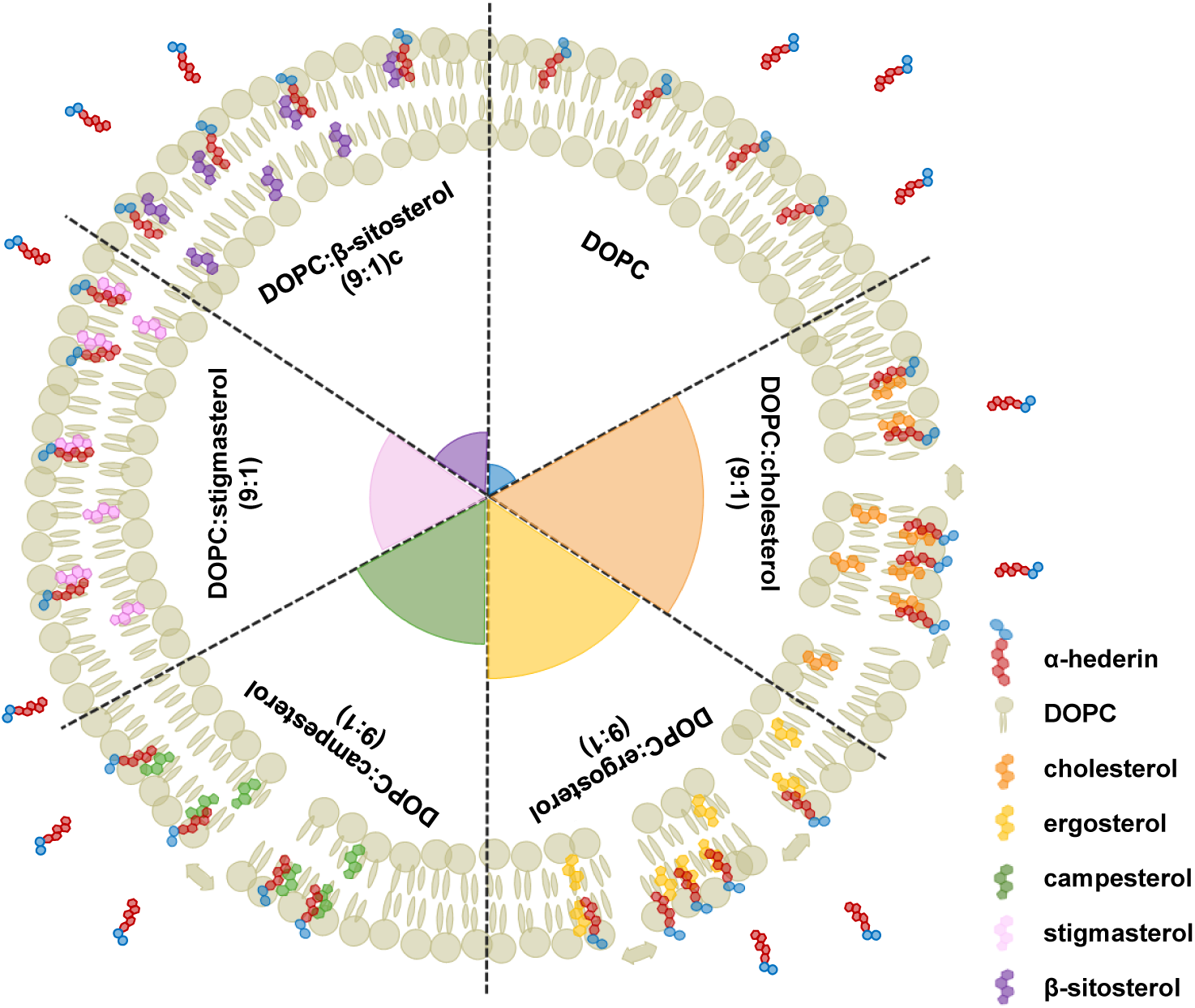
Model of saponin-sterol membrane interaction. Saponins induce membrane damage by increasing surface tension in specific regions through the formation of dense sterol–saponin aggregates. This aggregation is driven by the internal physicochemical properties of membranes containing animal or fungal sterols, rather than by species-specific binding affinity, which appears relatively uniform. In contrast, membranes rich in phytosterols do not promote such dense complex formation, resulting in reduced membrane disruption.

## REFERENCES

1. Espenshade, P. J. & Hughes, A. L. Regulation of sterol synthesis in eukaryotes. Annu Rev Genet 41, 401–427 (2007).

2. Brocks, J. J. et al. Lost world of complex life and the late rise of the eukaryotic crown. Nature 618, 767–773 (2023).

3. Sonawane, P. D. et al. Plant cholesterol biosynthetic pathway overlaps with phytosterol metabolism. Nat Plants 3, (2016).

4. Cárdenas, P. D. et al. Pathways to defense metabolites and evading fruit bitterness in genus Solanum evolved through 2-oxoglutarate-dependent dioxygenases. Nat Commun 10, 1–13 (2019).

5. Francis, G., Kerem, Z., Makkar, H. P. S. & Becker, K. The biological action of saponins in animal systems: a review. British Journal of Nutrition 88, 587–605 (2002).

6. Rai, S., Acharya-Siwakoti, E., Kafle, A., Devkota, H. P. & Bhattarai, A. Plant-Derived Saponins: A Review of Their Surfactant Properties and Applications. Sci 3, 1–19 (2021).

7. Singh, B. & Kaur, A. Control of insect pests in crop plants and stored food grains using plant saponins: A review. Lwt-Food Science and Technology 87, 93–101 (2018).

8. Wang, P. Natural and synthetic saponins as vaccine adjuvants. Vaccines (Basel) 9, 1–18 (2021).

9. Thimmappa, R., Geisler, K., Louveau, T., O’Maille, P. & Osbourn, A. Triterpene biosynthesis in plants. Annu Rev Plant Biol 65, 225–257 (2014).

10. Xu, P. & Yu, B. Chemical synthesis of saponins: An update. Advances in Carbohydrate Chemistry and Biochemistry, 1–62 (2021). doi:10.1016/bs.accb.2021.11.001.

11. Xu, R., Fazio, G. C. & Matsuda, S. P. T. On the origins of triterpenoid skeletal diversity. Phytochemistry 65, 261–291 (2004).

12. Augustin, J. M., Kuzina, V., Andersen, S. B. & Bak, S. Molecular activities, biosynthesis and evolution of triterpenoid saponins. Phytochemistry 72, 435–457 (2011).

13. Vincken, J. P., Heng, L., de Groot, A. & Gruppen, H. Saponins, classification and occurrence in the plant kingdom. Phytochemistry 68, 275–297 (2007).

14. da Silva Magedans, Y. V., Phillips, M. A. & Fett-Neto, A. G. Production of plant bioactive triterpenoid saponins: from metabolites to genes and back. Phytochemistry Reviews 20, 461–482 (2021).

15. Moses, T., Papadopoulou, K. K. & Osbourn, A. Metabolic and functional diversity of saponins, biosynthetic intermediates and semi-synthetic derivatives. Crit Rev Biochem Mol Biol 49, 439–462 (2014).

16. Lorent, J., Le Duff, C. S., Quetin-Leclercq, J. & Mingeot-Leclercq, M. P. Induction of highly curved structures in relation to membrane permeabilization and budding by the triterpenoid saponins, α- And δ-hederin. Journal of Biological Chemistry 288, 14000–14017 (2013).

17. Verstraeten, S. L. et al. The activity of the saponin ginsenoside Rh2 is enhanced by the interaction with membrane sphingomyelin but depressed by cholesterol. Sci Rep 9, 1–14 (2019).

18. Jacquier, N. & Schneiter, R. Mechanisms of sterol uptake and transport in yeast. J Steroid Biochem Mol Biol 129, 70–78 (2012).

19. Lorent, J. H., Quetin-Leclercq, J. & Mingeot-Leclercq, M. P. The amphiphilic nature of saponins and their effects on artificial and biological membranes and potential consequences for red blood and cancer cells. Org Biomol Chem 12, 8803–8822 (2014).

20. Hollingsworth, S. A. & Dror, R. O. Molecular Dynamics Simulation for All. Neuron 99, 1129–1143 (2018).

21. Li, J. & Monje-Galvan, V. Effect of Glycone Diversity on the Interaction of Triterpenoid Saponins and Lipid Bilayers. ACS Appl Bio Mater 7, 553–563 (2024).

22. Martinotti, C., Ruiz-Perez, L., Deplazes, E. & Mancera, R. L. Molecular Dynamics Simulation of Small Molecules Interacting with Biological Membranes. Chemphyschem 21, 1486–1514 (2020).

23. Bergman, L. W. Growth and Maintenance of Yeast BT - Two-hybrid systems — methods and protocols. Humana Press, 336 p., Two-hybrid systems — methods and protocols. Edited by Paul N. MacDonald, published by Humana Press, 2001, 336 p. 177, 9–14 (2001).

24. Sekula, B. C. & Nes, W. R. Metabolism of sterols by anaerobic *Saccharomyces cerevisiae*. Lipids 16, 195–198 (1981).

25. Achilles, J., Harms, H. & Müller, S. Analysis of living *S. cerevisiae* cell states - A three color approach. Cytometry Part A 69, 173–177 (2006).

26. Panaretou, B. & Piper, P. Isolation of yeast plasma membranes. Methods Mol Biol 313, 27–32 (2006).

27. Uzun, H.D., Vázquez-Hernández, M., Bandow, J. E. & Pomorski, T. G., *In vitro* Assay to Evaluate Cation Transport of Ionophores. Proteomics 22, 1–11 (2022).

28. Claereboudt, E. J. S., Eeckhaut, I., Lins, L. & Deleu, M. How different sterols contribute to saponin tolerant plasma membranes in sea cucumbers. Sci Rep 8, 1–11 (2018).

29. Quail, M. A. & Kelly, S. L. The Extraction and Analysis of Sterols from Yeast. in Yeast Protocols 123–132 (Humana Press, New Jersey). doi:10.1385/0-89603-319-8:123.

30. Khakimov, B., Motawia, M. S., Bak, S. & Engelsen, S. B. The use of trimethylsilyl cyanide derivatization for robust and broad-spectrum high-throughput gas chromatography-mass spectrometry based metabolomics. Anal Bioanal Chem 405, 9193–9205 (2013).

31. Trinh, M. D. L. et al. Site-directed genotype screening for elimination of antinutritional saponins in quinoa seeds identifies TSARL1 as a master controller of saponin biosynthesis selectively in seeds. Plant Biotechnol J 22, 2216–2234 (2024).

32. Jo, S., Kim, T., Iyer, V. G. & Im, W. CHARMM-GUI: A web-based graphical user interface for CHARMM. J Comput Chem 29, 1859–1865 (2008).

33. Jo, S., Lim, J. B., Klauda, J. B. & Im, W. CHARMM-GUI Membrane Builder for Mixed Bilayers and Its Application to Yeast Membranes. Biophys J 97, 50–58 (2009).

34. Wu, E. L. et al. CHARMM-GUI Membrane Builder toward realistic biological membrane simulations. J Comput Chem 35, 1997–2004 (2014).

35. Lee, J. et al. CHARMM-GUI Input Generator for NAMD, GROMACS, AMBER, OpenMM, and CHARMM/OpenMM Simulations Using the CHARMM36 Additive Force Field. J Chem Theory Comput 12, 405–413 (2016).

36. Hanwell, M. D., et al. Avogadro: an advanced semantic chemical editor, visualization, and analysis platform. J Cheminform 4, 17 (2012).

37. Kim, S. et al. CHARMM-GUI ligand reader and modeler for CHARMM force field generation of small molecules. J Comput Chem 38, 1879–1886 (2017).

38. Vanommeslaeghe, K. et al. CHARMM general force field: A force field for drug-like molecules compatible with the CHARMM all-atom additive biological force fields. J Comput Chem 31, 671–690 (2010).

39. Grubmüller, H., Heymann, B. & Tavan, P. Ligand binding: molecular mechanics calculation of the streptavidin-biotin rupture force. Science (1979) 271, 997–999 (1996).

40. Jarzynski, C. Nonequilibrium Equality for Free Energy Differences. Phys Rev Lett 78, 2690–2693 (1997).

41. Bonomi, M. et al. PLUMED: A portable plugin for free-energy calculations with molecular dynamics. Comput Phys Commun 180, 1961–1972 (2009).

42. Bonomi, M. et al. Promoting transparency and reproducibility in enhanced molecular simulations. Nat Methods 16, 670–673 (2019).

43. Tribello, G. A., Bonomi, M., Branduardi, D., Camilloni, C. & Bussi, G. PLUMED 2: New feathers for an old bird. Comput Phys Commun 185, 604–613 (2014).

44. Brooks, B. R. et al. CHARMM: The biomolecular simulation program. J Comput Chem 30, 1545– 1614 (2009).

45. Huang, J. et al. CHARMM36m: an improved force field for folded and intrinsically disordered proteins. Nat Methods 14, 71–73 (2017).

46. Mark, P. & Nilsson, L. Structure and Dynamics of the TIP3P, SPC, and SPC/E Water Models at 298 K. J Phys Chem A 105, 9954–9960 (2001).

47. Evans, D. J. & Holian, B. L. The Nose–Hoover thermostat. J Chem Phys 83, 4069–4074 (1985).

48. Parrinello, M. & Rahman, A. Polymorphic transitions in single crystals: A new molecular dynamics method. J Appl Phys 52, 7182–7190 (1981).

49. Abraham, M. J. et al. GROMACS: High performance molecular simulations through multi-level parallelism from laptops to supercomputers. SoftwareX 1, 19–25 (2015).

50. Hess, B., Bekker, H., Berendsen, H. J. C. & Fraaije, J. G. E. M. LINCS: A linear constraint solver for molecular simulations. J Comput Chem 18, 1463–1472 (1997).

51. Darden, T., York, D. & Pedersen, L. Particle mesh Ewald: An N⋅log(N) method for Ewald sums in large systems. J Chem Phys 98, 10089–10092 (1993).

52. Steinbach, P. J. & Brooks, B. R. New spherical-cutoff methods for long-range forces in macromolecular simulation. J Comput Chem 15, 667–683 (1994).

53. Humphrey, W., Dalke, A. & Schulten, K. VMD: Visual molecular dynamics. J Mol Graph 14, 33–38 (1996).

54. Michaud-Agrawal, N., Denning, E. J., Woolf, T. B. & Beckstein, O. MDAnalysis: A toolkit for the analysis of molecular dynamics simulations. J Comput Chem 32, 2319–2327 (2011).

55. Wang, C., et al. Glycosylation Drives Modulation of Triterpenoid Saponin Solubility, Micelle Formation, and Stability. (2025) 10.26434/chemrxiv-2025-28qzd.

56. Hillenmeyer, M. E. et al. The Chemical Genomic Portrait of Yeast: Uncovering a Phenotype for All Genes. Science (1979) 320, 362–365 (2008).

57. Johnston, E. J. et al. Yeast lacking the sterol C-5 desaturase Erg3 are tolerant to the anti-inflammatory triterpenoid saponin escin. Sci Rep 13, 1–11 (2023).

58. Wang, W. et al. Functional Analysis of the C-5 Sterol Desaturase PcErg3 in the Sterol Auxotrophic Oomycete Pathogen Phytophthora capsici. Front Microbiol 13, 1–11 (2022).

59. Lorent, J. et al. Domain formation and permeabilization induced by the saponin α-hederin and its aglycone hederagenin in a cholesterol-containing bilayer. Langmuir 30, 4556–4569 (2014).

60. Lin, F. & Wang, R. Hemolytic mechanism of dioscin proposed by molecular dynamics simulations. J Mol Model 16, 107–118 (2010).

61. Thimmappa, R. et al. Biosynthesis of saponin defensive compounds in sea cucumbers. Nature Chemical Biology 18, 774–781 (2022).

62. Yue, Z., Li, C., Voth, G. A. & Swanson, J. M. J. Dynamic Protonation Dramatically Affects the Membrane Permeability of Drug-like Molecules. J Am Chem Soc 141, 13421–13433 (2019).

63. Kuljanabhagavad, T., Thongphasuk, P., Chamulitrat, W. & Wink, M. Triterpene saponins from Chenopodium quinoa Willd. Phytochemistry 69, 1919–1926 (2008).

64. Kuljanabhagavad, T. & Wink, M. Biological activities and chemistry of saponins from *Chenopodium quinoa* Willd. Phytochemistry Reviews 8, 473–490 (2009).

65. Lacchini, E. et al. The saponin bomb: a nucleolar-localized β-glucosidase hydrolyzes triterpene saponins in *Medicago truncatula*. New Phytologist, 239(2), 705–719 (2023).

