## Supplemental Figures S1-S4 for "Sterols govern membrane susceptibility to saponin-induced lysis"

### Supplemental Figures S1-S4
### Sterols govern membrane susceptibility to saponin-induced lysis


Malbor Dervishi[1], Jan Günther[1], Jinhui Li[2], Huriye Deniz Uzun[1,3], Hans Christian Bruun Hansen[1], Thomas Gümther-Pomorski[1,3], Anja Thoe Fuglsang[1], Viviana Monje[2], and Søren Bak[1,*]

*Department of Plant and Environmental Sciences, University of Copenhagen, Thorvaldsensvej 40, 1871 Frederiksberg, Denmark[1]*

*Department of Chemical and Biological Engineering, University of Buffalo, 308 Furnas Hall, Amherst, NY 14260, United States[2]*

*Department of Molecular Biochemistry, Faculty of Chemistry and Biochemistry, Ruhr University Bochum, 44780 Bochum, Germany[3]*

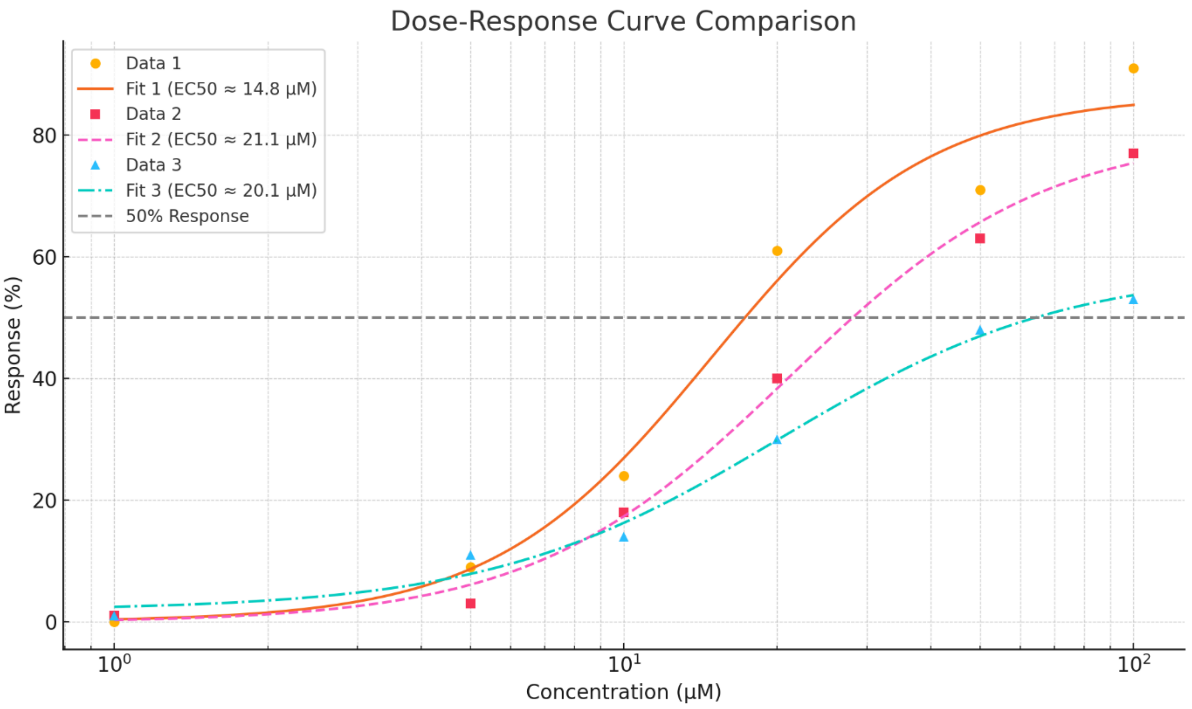


**Figure S1:** Dose response curves for LUVs composed of DOPC:CHOL (Data 1), DOPC:ERGO (Data 2) and DOPC:CAMP (Data 3).

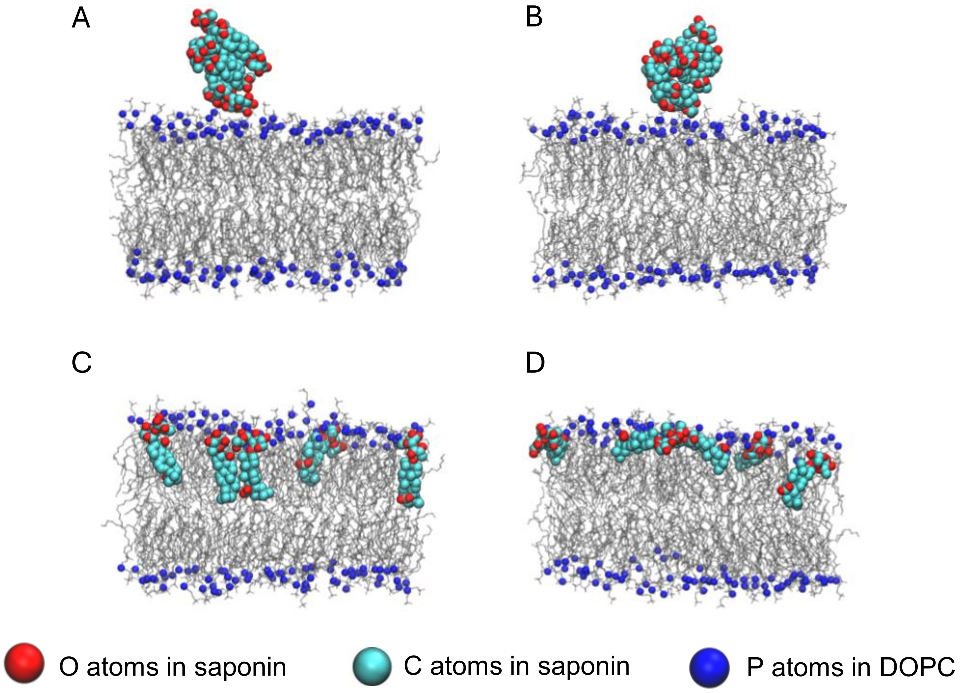


**Figure S2:** Final stages of the simulation systems with five α-hederin molecules initially located in the water phase near the bilayer for the (A) neutral and (B) deprotonated form and five α-hederin molecules initially in an embedded conformation in the DOPC/CHOL bilayer model for the (C) neutral and (D) deprotonated forms.

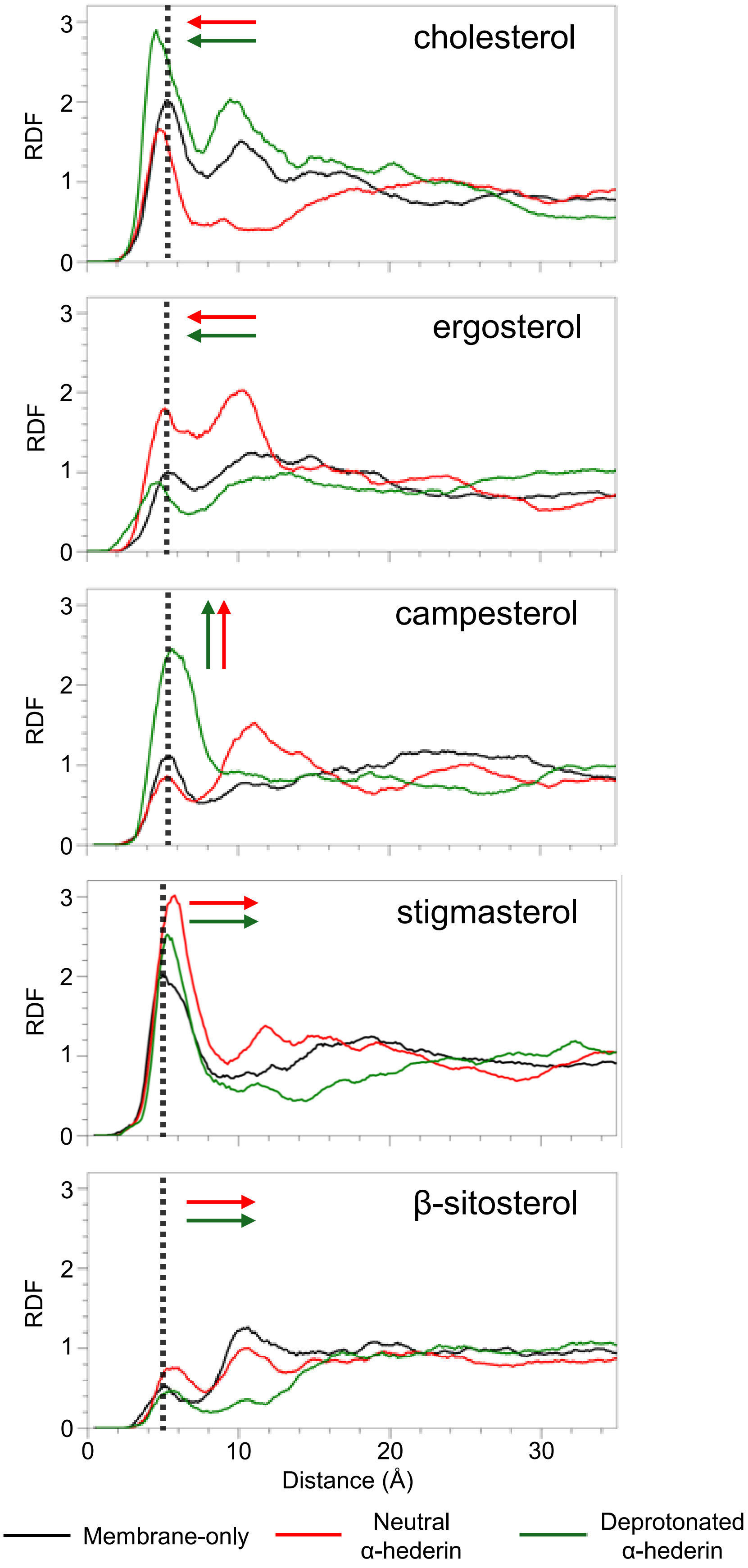


**Figure S3:** Two-dimensional radial distribution functions (RDFs) of sterol–sterol interactions. The curves represent the probability distribution of finding sterol molecules near each other, calculated based on the lateral (x–y plane) distance between their hydroxyl oxygen atoms. Black, red, and green lines correspond to membrane-only systems, membranes containing five neutral α-hederin molecules, and membranes with five deprotonated α-hederin molecules, respectively. Vertical grey dashed lines indicate the position of the first solvation shell in membrane-only systems (i.e., without saponins).

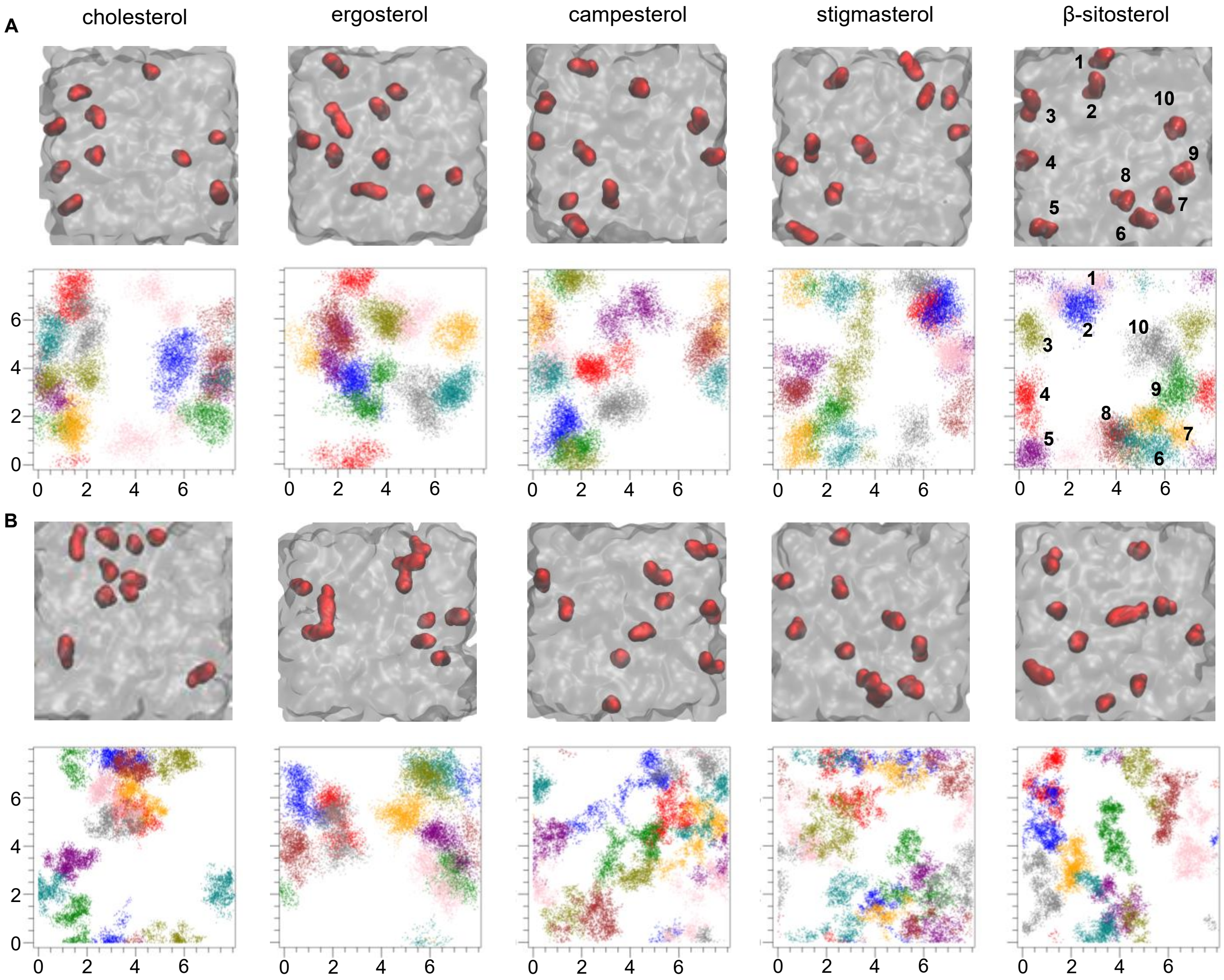


**Figure S4: Molecular modeling of membrane-saponin interaction provides mechanistic insights into saponin bioactivity.** Representative top-view snapshots and corresponding cumulative sterol distribution maps from the last 100 ns of simulations for (A) membranes-only systems and (B) membranes with five bound deprotonated α-hederin molecules. Red globes represent sterols in the α-hederin-bound leaflet; grey represents DOPC lipids; and colorful scatter points in the distribution map represent the positions of the individual sterol hydroxyl oxygen atoms. Individual cholesterol molecules are numbered in the last column as reference. We observe a higher natural aggregation of the sterols compared to the phytosterols. The aggregation of CHOL and ERGO increases with the presence of saponins.
