## Supplemental Tables S1-S3 for "Sterols govern membrane susceptibility to saponin-induced lysis"

### Sterols govern membrane susceptibility to saponin-induced lysis

Malbor Dervishi[1], Jan Günther[1], Jinhui Li[2], Huriye Deniz Uzun[1,3], Hans Christian Bruun Hansen[1], Thomas Gümther-Pomorski[1,3], Anja Thoe Fuglsang[1], Viviana Monje[2], and Søren Bak[1,*]

*Department of Plant and Environmental Sciences, University of Copenhagen, Thorvaldsensvej 40, 1871 Frederiksberg, Denmark[1]*

*Department of Chemical and Biological Engineering, University of Buffalo, 308 Furnas Hall, Amherst, NY 14260, United States[2]*

*Department of Molecular Biochemistry, Faculty of Chemistry and Biochemistry, Ruhr University Bochum, 44780 Bochum, Germany[3]*

**Table S1.** Summary of saponin-in-water and DOPC/sterol membrane-only systems.

| System Names | Lipids /leaflet | Total atoms # | Simulation box dimensions (nm) | Sim. time × reps (ns) |
|---|---|---|---|---|
| **Saponin-in-water** | | | | |
| Neutral α-hederin | - | 6989 | 4.11*4.11*4.11 | 150×3 |
| Deprotonated α-hederin | - | 6983 | 4.11*4.11*4.11 | 150×3 |
| hederacoside C | - | 6969 | 4.11*4.11*4.11 | 150×3 |
| **Pure DOPC/sterol membranes** | | | | |
| DOPC/CHOL | 100 | 62433 | 7.78*7.78*9.93 | 300×3 |
| DOPC/ERGO | 100 | 62353 | 7.99*7.99*9.44 | 300×3 |
| DOPC/CAMP | 100 | 62478 | 7.88*7.88*9.73 | 300×3 |
| DOPC/STIG | 100 | 62444 | 8.04*8.04*9.33 | 300×3 |
| DOPC/ β-SITO | 100 | 62529 | 7.91*7.91*9.92 | 300×3 |

**Table S2.** Summary of simulated systems with protonated α-hederin (AHD), deprotonated α-hederin (AHN), and hederacoside C (HDC), respectively.

| Saponin | Sterol | Lipids /leaflet | Total atoms # | Simulation box dimensions (nm) | Sim. time × reps (ns) |
|---|---|---|---|---|---|
| **Single saponin-membrane (saponin starts from water)** | | | | | |
| AHD | CHOL | 100 | 59823 | 7.91*7.91*9.22 | 500×3 |
| | ERGO | 100 | 59623 | 7.83*7.83*9.33 | 500×3 |
| | CAMP | 100 | 59883 | 7.82*7.82*9.41 | 500×3 |
| | STIG | 100 | 59323 | 7.74*7.74*9.54 | 500×3 |
| | β-SITO | 100 | 60050 | 7.89*7.89*9.26 | 500×3 |
| AHN | CHOL | 100 | 59817 | 7.76*7.76*9.52 | 500×3 |
| | ERGO | 100 | 59605 | 7.96*7.96*9.03 | 500×3 |
| | CAMP | 100 | 59877 | 7.90*7.90*9.26 | 500×3 |
| | STIG | 100 | 59329 | 7.82*7.82*9.32 | 500×3 |
| | β- SITO | 100 | 60056 | 7.91*7.91*9.18 | 500×3 |
| HDC | CHOL | 100 | 59816 | 7.83*7.83*9.39 | 500×3 |
| | ERGO | 100 | 59637 | 7.88*7.88*9.24 | 500×3 |
| | CAMP | 100 | 59882 | 7.78*7.78*9.52 | 500×3 |
| | STIG | 100 | 59319 | 7.86*7.86*9.25 | 500×3 |
| | β-SITO | 100 | 60037 | 7.88*7.88*9.29 | 500×3 |
| **5 saponin-membrane (Saponins start from water)** | | | | | |
| 5xAHD | CHOL | 100 | 59777 | 7.73*7.73*9.54 | 500 |
| | ERGO | 100 | 59580 | 7.93*7.93*9.11 | 500 |
| | CAMP | 100 | 59816 | 7.89*7.89*9.22 | 500 |
| | STIG | 100 | 59325 | 7.83*7.83*9.28 | 500 |
| | β-SITO | 100 | 60004 | 7.94*7.94*9.12 | 500 |
| 5xAHN | CHOL | 100 | 59738 | 7.94*7.94*9.05 | 500 |
| | ERGO | 100 | 59559 | 7.77*7.77*9.47 | 500 |
| | CAMP | 100 | 59789 | 7.82*7.82*9.36 | 500 |
| | STIG | 100 | 59265 | 7.91*7.91*9.09 | 500 |
| | β-SITO | 100 | 59959 | 7.88*7.88*9.23 | 500 |
| **5 saponin-membrane (Saponins start from membrane core region)** | | | | | |
| 5xAHD | CHOL | 100 | 59777 | 8.01*8.01*8.91 | 500×3 |
| | ERGO | 100 | 59580 | 7.97*7.97*9.01 | 500×3 |
| | CAMP | 100 | 59816 | 7.95*7.95*9.05 | 500×3 |
| | STIG | 100 | 59325 | 7.98*7.98*8.94 | 500×3 |
| | β-SITO | 100 | 60004 | 7.83*7.83*9.36 | 500×3 |
| 5xAHN | CHOL | 100 | 59738 | 7.98*7.98*8.99 | 500×3 |
| | ERGO | 100 | 59559 | 8.08*8.08*8.78 | 500×3 |
| | CAMP | 100 | 59789 | 8.01*8.01*8.93 | 500×3 |
| | STIG | 100 | 59265 | 7.92*7.92*9.03 | 500×3 |
| | β-SITO | 100 | 59959 | 8.05*8.05*8.86 | 500×3 |

**Table S3**. Summary of stable binding events for neutral α-hederin (AHD), deprotonated α-hederin (AHN), and hederacoside C (HDC) in single saponin–membrane systems. Values are reported as binding events/total simulations.

| | AHD | AHN | HDC |
|---|---|---|---|
| DOPC/CHOL | 1/3 | 2/3 | 1/3 |
| DOPC/ERG | 1/3 | 3/3 | 2/3 |
| DOPC/CAMP | 2/3 | 1/3 | 1/3 |
| DOPC/SITG | 3/3 | 0/3 | 0/3 |
| DOPC/β-SITO | 3/3 | 2/3 | 0/3 |
| Total | 10/15 | 8/15 | 4/15 |
